# Fatty acid synthase coordinates mTOR signaling and histone crotonylation during cardiomyocyte hypertrophy

**DOI:** 10.64898/2026.10.26.762072

**Authors:** Angeliki Ntorla, Pierre Coleman, Yujing Xia, Helen R. Warren, Joseph Burgoyne

## Abstract

**Background:** Epigenetic modifications are central to the regulation of cardiac gene expression during disease. Among these, histone crotonylation (Kcr) has recently emerged as a key modulator of chromatin structure and transcriptional activity. Here, we identify a novel signaling axis linking fatty acid synthase (FASN) to mTOR activation and hypertrophic gene expression through histone crotonylation.

**Methods:** FASN expression, mTOR activation, and histone crotonylation were assessed in mouse hearts and isolated cardiomyocytes using immunoblotting. FASN was silenced by siRNA or inhibited pharmacologically using TVB-2640. Hypertrophic gene expression was quantified by RT-qPCR. Cardiomyocyte hypertrophy was evaluated by immunofluorescence-based cell size analysis. p300 acetylation and crotonylation were assessed by immunoprecipitation and immunoblotting. CUT&RUN-seq and RNA-seq were performed to identify FASN- and Histone H3K18cr-regulated genes during hypertrophic stimulation. Bioinformatic analyses of human RNA-seq and genome-wide association study (GWAS) data were conducted to investigate the clinical relevance of these findings.

**Results:** Histone crotonylation was markedly elevated in hypertrophic myocardium and cardiomyocytes. This increase was driven by upregulated FASN expression in response to hypertrophic stimuli. Genetic or pharmacological inhibition of FASN suppressed histone crotonylation, hypertrophic gene expression, and cardiomyocyte enlargement. Mechanistic studies suggested that FASN regulates histone crotonylation through mTOR-associated signaling and changes in p300 activity. Transcriptomic analyses further revealed that FASN enhances the expression of hypertrophy-associated genes enriched for histone crotonylation marks. Analysis of human sequencing datasets demonstrated a positive association between FASN expression and molecular markers of hypertrophy in patients with hypertrophic cardiomyopathy (HCM). Furthermore, GWAS analysis identified a variant at the FASN locus associated with heart failure as the underlying cause of death and increased FASN expression in cardiomyocytes.

**Conclusions:** Our findings identify FASN as an important regulator of mTOR signalling and histone crotonylation during cardiomyocyte hypertrophy. These results support a potential mechanism linking altered lipid metabolism to epigenetic regulation and pathological cardiac remodelling.

## Introduction

Heart failure is a complex clinical syndrome characterized by the inability of the heart to pump sufficient blood to meet the metabolic demands of the body and remains a leading cause of morbidity and mortality worldwide. While traditional research has emphasized genetic mutations and hemodynamic stress as primary drivers of cardiac dysfunction, growing evidence indicates that epigenetic regulation plays a critical role in the initiation and progression of heart failure^1^. Epigenetic mechanisms, including DNA methylation, histone modifications, and non-coding RNAs, modulate gene expression without altering the underlying DNA sequence, enabling cardiac cells to dynamically respond to environmental, metabolic, and mechanical stressors^2^. These regulatory processes influence key pathological features such as myocardial hypertrophy, fibrosis, inflammation, metabolic remodeling, and cardiomyocyte survival^1,3^.

Histone lysine crotonylation (Kcr) is a newly identified epigenetic modification that has a pronounced capacity to regulate gene expression^4,5^. This modification has been implicated in HIV latency^6^, acute kidney injury^7^, and cancer^8^. In addition, elevation in histone crotonylation has been associated with cardiac hypertrophy^9^, where it is upregulated in human hearts with hypertrophic cardiomyopathy (HCM) and may promote recruitment of the NFATC3 transcription factor to the *Nppb* promoter. It has been proposed that this increase in histone Kcr results from reduced expression of ECHS1, an enzyme that catalyses hydration of crotonyl-CoA into (S)-3-hydroxybutyryl-CoA^10^.

In the present study, we sought to more precisely define the role of histone Kcr in cardiomyocyte hypertrophy and to elucidate the mechanisms regulating this modification. Under our experimental conditions we did not observe evidence that reduced ECHS1 expression accounts for the increase in histone crotonylation^9^. Instead, our findings implicate increased fatty acid synthase (FASN) expression in the regulation of histone crotonylation during cardiomyocyte hypertrophy. We further identify an association between FASN-dependent mTOR signalling, changes in p300 activity and increased histone H3K18cr.

Finally, transcriptomic and epigenomic analyses suggest that histone crotonylation contributes to the regulation of genes associated with cardiomyocyte hypertrophy and cardiomyopathy.

## Methods

### Cell culture

Primary culture of neonatal rat ventricular myocytes (NRVMs) was prepared by adapting previously described protocols^11,12^. Briefly, after sacrifice of 1- to 3-day-old Wistar rat pups, hearts were immediately collected and transferred into ice-cold ADS buffer (116 mM NaCl, 20 mM HEPES pH 7.35, 1 mM NaH_2_PO_4_, 5.6 mM glucose, 5.4 mM KCl, 0.8 mM MgSO_4_).

After the removal of lung tissue, large vessels and atria, hearts were rinsed in the ADS buffer and minced into small pieces (about 1mm^3^). Next, ventricle pieces were completely dissociated by repeated enzymatic digestion with 0.8 mg/mL collagenase type 2 (Worthington Biochemical, cat. #LS004176) and 0.6 mg/mL pancreatin (Sigma-Aldrich, cat. #P3292) in ADS buffer. Undigested tissue was cleared by cell strainer and cells were recovered by centrifugation at 1250 rpm for 5 min. The cell pellet was resuspended in plating medium [4:1 mixture of DMEM (Sigma-Aldrich, cat. #D-6546) and Medium 199 (M199) (Sigma-Aldrich, cat. #M-2154) supplemented with 10% horse serum (Sigma-Aldrich, cat. #H1270), 5% non-heat inactivated FBS (Sigma-Aldrich, cat. #F7524), 1% glutamine/P/S (Sigma-Aldrich, cat. #G1146), 1% non-essential amino acids (Gibco™, cat. #11140050)] and preplated in T175 flasks for 1 hour to eliminate easy-adherent non-cardiomyocytes. Finally, unattached cardiomyocytes were harvested and plated on gelatin-coated (Sigma-Aldrich, cat. #G1393) 6-well or 12-well plates at 250,000 cells/mL. After 18 h, neonatal cardiomyocytes were washed and cultured in serum-free maintenance medium [4:1 mixture of DMEM and M199 supplemented with 1% glutamine/P/S and 0.1 mM 5-bromo-2’-deoxyuridine (Sigma-Aldrich, cat. #B5002)] for the duration of the experiments. Cells were maintained at 37°C in a 5% CO_2_ humidified incubator and were treated with vehicle, endothelin-1 (ET-1; Sigma-Aldrich, cat. #E7764), phenylephrine (PE; Sigma-Aldrich, cat. #P6126), angiotensin-II (AngII; Sigma-Aldrich, cat. #A9525), rapamycin (Cell Signaling Technology, cat. #9904) or TVB-2640 (Selleck, cat. #S9714)

### Western blot

Protein extracts were prepared in sample buffer containing 50 mM Tris-HCl pH 6.8, 2% w/v sodium dodecyl sulfate, 10% glycerol, 0.0025% w/v bromophenol blue and 5% β-mercaptoethanol. Samples were heated for 5 min at 95°C before being resolved on 4-15% Mini-PROTEAN® TGX™ Gels (BioRad, cat. #4561086) and then transferred onto methanol activated polyvinylidene fluoride (PVDF) membranes (BioRad) using a Trans-Blot Turbo Transfer System (BioRad). The PVDF membranes were then blocked with 10 % non-fat dry milk or 10% bovine serum albumin (BSA) (Sigma-Aldrich, cat. #A9647) in phosphate buffered saline (PBS) (Seven Biotech Ltd., cat. #20-7461-01) with 0.1 % v/v Tween® 20 (VWR International, cat. #663684B) (PBS-T) for 1 hour, and then incubated with the primary antibodies for 2 hours at room temperature or overnight at 4°C. Following that, they were washed 3 times with PBS-T, incubated with the appropriate secondary antibody for 1 hour, washed again and developed with the Pierce ECL detection system (Thermo-Fisher Scientific, cat. #32106). In a dark room, membranes were exposed to photographic film (GE Healthcare, cat. #28-9068-37) and chemiluminescent signal was detected using an automatic processor (Fuji RG II). Western blot quantification was performed using the Gel-Pro Analyser (version 3.1). Antibodies used in this study include PanKcr (PTM Biolabs, PTM-509), H3K9cr (PTM Biolabs, PTM-516), H3K18cr (Abcam, ab195475), PanKAc (Cell Signaling Technology, 9441), anti-H3 (Cell Signaling Technology, 4499), ECHS1 (Atlas Antibodies, HPA022476), FASN (Cell Signaling Technology, 3180), Phosphop70 S6 Kinase (Thr389, Cell Signaling Technology, 9205), p70 S6 Kinase (Cell Signaling Technology, 9202), Alpha Skeletal Muscle Actin (Abcam, ab28052), ANP (Proteintech, 27426-1-AP), GAPDH (Cell Signaling Technology, 2118), β-Actin (Merck, A5316), Anti-rabbit IgG, HRP-linked secondary antibody (Cell Signaling Technology, 7074), Anti-mouse IgG, HRP-linked secondary antibody (Cell Signaling Technology, 7076).

### Immunoprecipitation

For p300 immunoprecipitation, cells were lysed in ice-cold lysis buffer from Cell Signaling Technology (cat. #9803) supplemented with a complete protease inhibitor cocktail (Roche, 05892791001) and 5mM sodium butyrate. Samples were incubated with p300 primary antibody (Cell Signaling Technology, 86377) overnight at 4°C, followed by the addition of pre-washed protein A magnetics beads (Cell Signaling Technology, cat. #73778).

Immunocomplexes were washed extensively five times with lysis buffer and then subjected to Western blotting.

### Immunofluorescence staining

NRVMs were fixed in ice-cold methanol for 15 minutes. After 3 washes with PBS, samples were blocked with 5% BSA in PBS for 1 hour at room temperature and then incubated with the appropriate dilution of primary antibody (1:1000, troponin, Abcam ab56357) overnight at 4°C in PBS. Following 3 washes with PBS, samples were incubated with Alexa Fluor 488 secondary antibody (Invitrogen) for 1 hour at room temperature and nuclei were counterstained with Hoechst 33342 (Invitrogen, cat. #H3570) for 10 minutes. The stained cells were analysed by an EVOS microscope (ThermoFisher Scientific) and cell size was measured by ImageJ.

### Animal model of pressure overload induced hypertrophy

Animal experiments were carried out by Dr Brian Cooley at the University of North Carolina School of Medicine. In brief, 9-week old male C57Bl/6 mice purchased from The Jackson Laboratory underwent transverse aortic constriction (TAC). Mice were anaesthetised with a mixture of 100 mg/kg ketamine and 15 mg/kg xylazine via IP injection. Aortic arch was accessed via the suprasternal notch approach with a 2-3 mm sternal cut. Trans-aortic arch constriction was achieved by ligation of the aorta between innominate and left carotid arteries with 6-0 silk, using 27 1⁄2 gauge spacer. For Sham mice, these underwent the same surgical operation but without the constrictive suture. 7 days after the operation mice were culled and hearts were snap-frozen in liquid nitrogen and stored at -80°C until processing. For immunoblot analysis hearts were ground to a fine powder using a pestle and mortar under liquid nitrogen, with a portion then added to sample buffer.

### siRNA transfections

siRNA transfections were performed using Lipofectamine™ RNAiMAX (Invitrogen, cat. #13778030) as per manufacturer’s instruction. The following siRNAs were used at the following concentrations: non-target control pool (Dharmacon D-001810-10-05, 50 nM), Fasn #1 (Dharmacon J-095791-09, 50 nM) or SMARTpool Echs1 (Dharmacon L-092231-02).

### qPCR analysis

Total RNA was extracted from cells using a RNeasy® Mini Kit (Qiagen, cat. #74104), following the instructions provided by the manufacturer. To eliminate genomic DNA contamination, on-column DNase digestion was performed using the RNAse-Free DNase Set (Qiagen, cat. #79254). RNA quantification and purity control were performed by measuring its absorbance at 260 nm and 280 nm with a NanoDrop™ 2000c Spectrophotometer (Thermo Fisher Scientific, cat. #ND200C).

Approximately 400 ng of total RNA was reverse transcribed to cDNA using the High-Capacity cDNA Reverse Transcription kit (Applied Biosystems™, cat. #4368814) following the manufacturer’s instructions. qPCR analysis was performed with the Luna Universal qPCR Master mix (New England Biolabs, cat. #M3003L) and primers for rat ANP mRNAs ( forward: 5’- CTGGGACCCCTCCGATAGAT-3’, reverse: 5’-CAATCCTACCCCCGAAGCAG-3’,) and FASN mRNAs (forward: 5’-TGTACCCTCTAGCTGGACCC-3’, reverse: 5’-CCAGGCTAAGGGCAATGGAA-3’) using the 7900HT Fast Real-Time PCR system (Applied Biosystems™). For each reaction, 2 μL cDNA was amplified using 5 μL of the SYBR Green I containing Luna master mix, 0.25 μL of both forward and reverse primer (10 μM) and nuclease-free water in a total reaction volume of 10 μL. All reactions were run in triplicates with the standard two-step cycling protocol.

### CUT&RUN assay and library preparation

The CUT&RUN experiment was carried out using the CUT&RUN Assay Kit from Cell Signaling Technology (cat. #86652) following the manufacturer’s protocol. Primary cardiomyocytes were harvested, counted, and centrifuged for 3 min at 600 x g at room temperature. 125,000 cells were used for each antibody reaction and IgG control sample. DNA from input and enriched chromatin samples was purified using the DNA Purification Buffers and Spin Columns kit for ChIP and CUT&RUN (Cell Signaling Technology, cat. #14209) and eluted in 40 μL Elution Buffer. CUT&RUN libraries were prepared using the NEBNext® Ultra™ II DNA Library Prep Kit for Illumina® (New England Biolabs, cat. #E7645) and NEBNext® Multiplex Oligos for Illumina® (Index Primers Set 1, cat. #7335) according to manufacturer’s instructions with minor modifications for CUT&RUN samples as recommended in the CUT&RUN kit protocol. Paired-end Illumina sequencing (2×150bp) was conducted on pooled libraries using the NovaSeq™ 6000 system by Novogene. Data analysis was performed by Novogene.

### RNA-seq analysis

RNA-seq libraries were prepared at Novogene by using the mRNA poly(A) selection method and sequenced on an Illumina NovaSeq platform (2×150bp). Downstream bioinformatics analysis was also carried out by Novogene. In brief, sequencing reads were processed with the fastp software to remove possible adapter sequences and low-quality nucleotides. The trimmed reads were then aligned to the rat genome (Rnor6.0) using the HISAT2. Unique gene hit counts were calculated using featureCounts. Differential gene expression was performed with DESeq2. Finally, gene ontology (GO) and KEGG enrichment analysis was performed on the significantly differentially expressed genes using the clusterProfiler.

### Human HCM analysis

RNAseq raw counts of cardiac tissue from 97 patients with symptomatic obstructive HCM were obtained from a study by Garmany et al^13^. Counts were processed with DESeq2^14^.

Genes were retained if they had at least 10 counts in at least 2 samples, and variance-stabilizing transformed (VST) expression values were generated for downstream correlation analyses. Gene-wise associations with FASN were then tested using ordinary least-squares regression. P-values were adjusted for multiple comparisons using the Benjamini-Hochberg method. For specified correlation analyses, VST-normalised expression values were residualised against the sample-wide mean expression. For every gene, ordinary least-squares regression was used to model expression as a function of the mean expression across all retained genes in each sample, and residuals from these models were used. Genes annotated to HCM were obtained from KEGG pathway hsa05410.

### Analysis of publicly available GWAS and eQTL data

A locus-zoom plot was created for variants associated with underlying (primary) cause of death: ICD10: I50.9 Heart failure (UK Biobank Data-Field 40001) within FASN gene locus using PheWeb^15^. The association of variants with FASN expression in iPSC-derived cardiomyocytes was assessed by analyzing eQTLs from the Elorbany et al. dataset^16^, using SingleQ^17^.

### Statistical analysis

The numbers of independent experiments are indicated in the relevant figure legends. Data are represented as mean and standard error of the mean (SEM). Statistical analyses were performed using GraphPad Prism software (v.8.4.3) unless stated otherwise. Normal distribution of data was examined by Shapiro-Wilk normality test. For statistical comparisons between two groups, unpaired two-tailed Student’s t-test was applied. Comparisons between 3 or more groups were conducted by one-way ANOVA followed by Holm-Sidak’s post hoc test or Kruskal-Wallis analysis followed by Dunn’s post-hoc test when data were not normally distributed. p-values <0.05 were considered statistically significant for all comparisons.

## Results

### Hypertrophic stimulation enhances histone crotonylation

In cardiomyocytes exposed to hypertrophic stimuli, an increase in both total and site-specific histone Kcr was observed (Fig. 1A). Similarly, total histone crotonylation was elevated, and a trend for enhanced site specific Kcr was shown, seven days post-transverse aortic constriction (TAC) in hypertrophic myocardium (Fig. 1B). These results align with previous findings, where elevated histone crotonylation was also observed in human hearts with HCM^9^.

**Figure 1.**
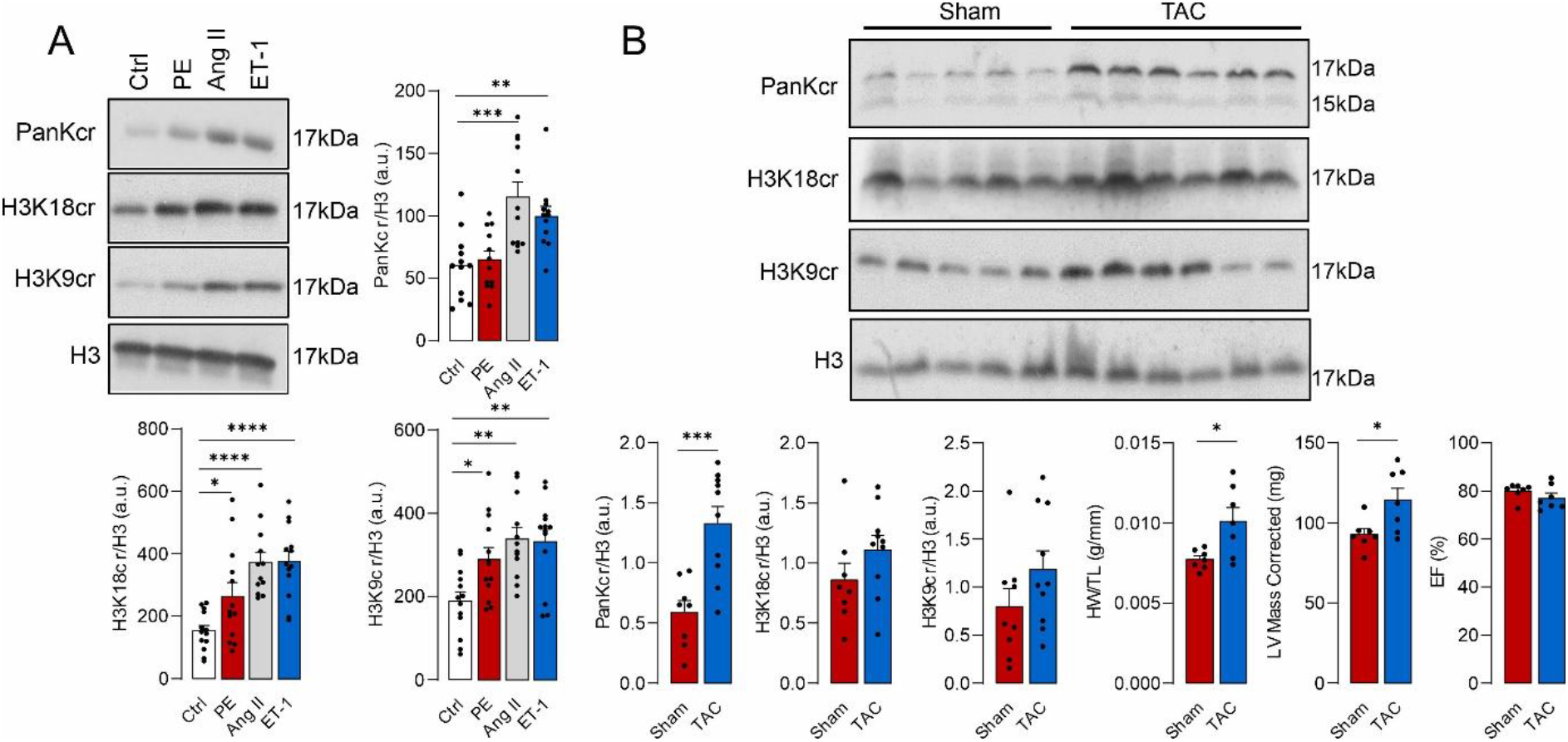
Histone crotonylation is elevated in cardiomyocytes and the myocardium with hypertrophic stimulation. **A**, Pan histone Kcr and site specific H3K9cr and H3K18cr in cardiomyocytes treated with hypertrophic stimuli for 48 hours (n=12-13). B, Pan histone Kcr and site specific H3K9cr and H3K18cr, as well as HW/TL, LV Mass and EF of hearts from mice 7-days post Sham or TAC surgery (n=7-10). *P<0.05; **P<0.01; ***P<0.001; ****P<0.0001. Comparisons were made using 1-way ANOVA followed by the Tukey post hoc test (**A**), or Student’s t-test (**B**).

### Elevation in FASN expression drives histone crotonylation and cardiomyocyte hypertrophy

Increased histone Kcr associated with cardiac hypertrophy has been attributed to the loss of ECHS1 during heart failure progression^9^. However, loss of ECHS1 under conditions of elevated histone Kcr were not observed when cardiomyocytes were treated with ET-1 or Ang-II, or in the myocardium post-TAC surgery (Fig. 2A, B). Additionally, knockdown (KD) of ECHS1 did not impact on total histone Kcr but led to a decrease in H3K18cr (Fig. S1).

**Figure 2.**
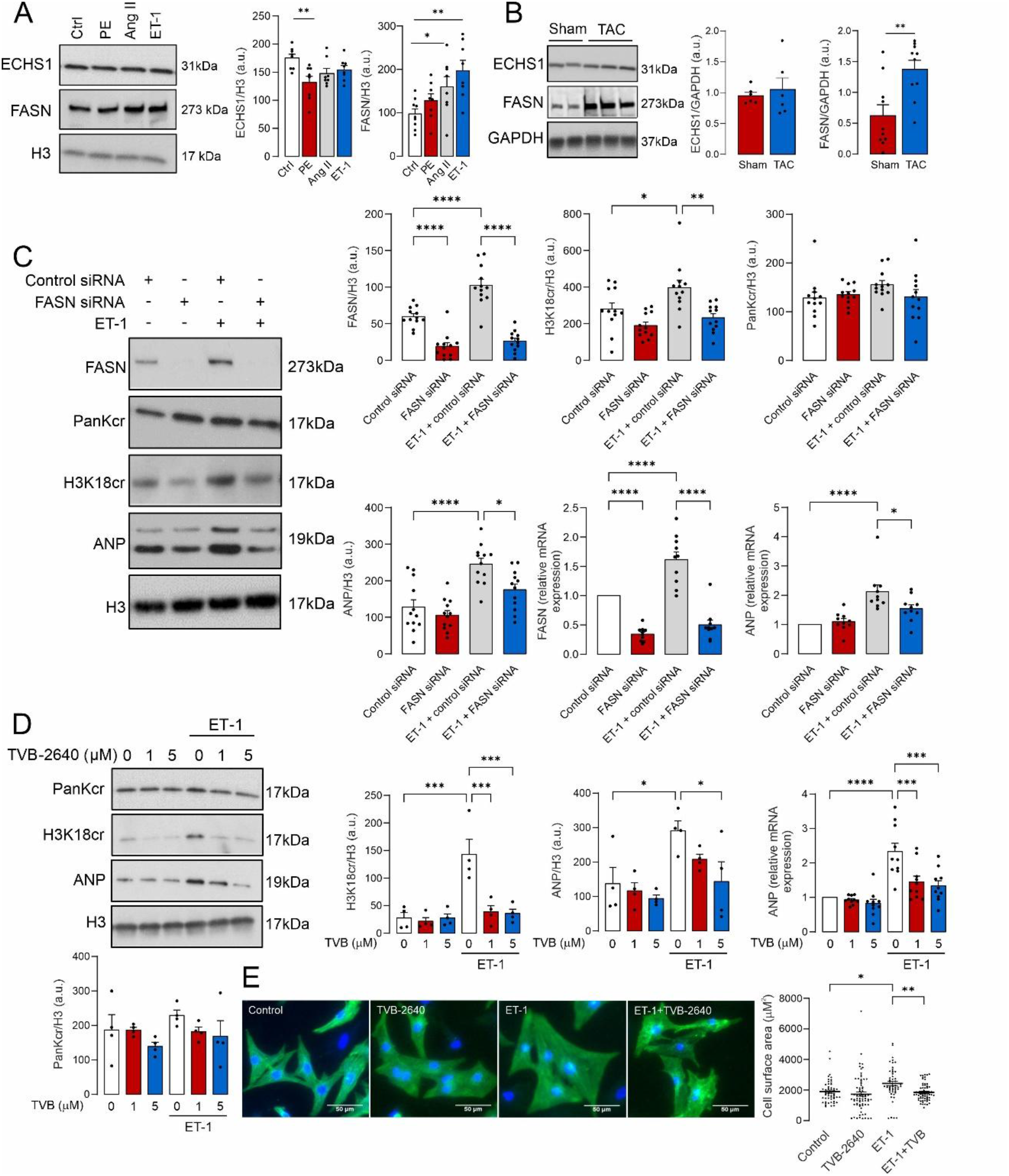
Upregulation of fatty acid synthase drives elevated histone crotonylation and cardiomyocyte hypertrophy. **A**, ECHS1 and FASN expression in cardiomyocytes treated with hypertrophic stimuli (n=8-9). **B**, ECHS1 and FASN expression in hearts from mice 7-days post Sham or TAC surgery (n=6, 10). **C**, Histone Kcr, H3K18cr, FASN and ANP expression in cardiomyocytes with or without knock-down of FASN and treated with or without ET-1 (n=12, 10). **D**, Histone Kcr, H3K18cr and ANP expression in cardiomyocytes treated with or without the FASN inhibitor TVB-2640 and in the presence or absence of ET-1 (n=4, 10). **E**, Cardiomyocyte surface area measured in cardiomyocytes treated with or without the FASN inhibitor TVB-2640 and in the presence or absence of ET-1. *P<0.05; **P<0.01; ***P<0.001; ****P<0.0001. Comparisons were made using 1-way ANOVA followed by the Tukey post hoc test (**A, C, D, E**), or Student’s t-test (**B**).

This result is not entirely unexpected, given crotonyl-CoA, the substrate for histone crotonylation, can also be generated by ECHS1^18,19^.

To investigate further, we examined other pathways that could influence crotonyl-CoA formation. This led us to hypothesize that upregulation of fatty acid synthase (FASN) seen in cardiomyocytes following hypertrophic stimulation and in hypertrophic myocardium (Fig. 2A, B) may contribute to increased histone crotonylation, due to its impact on cellular metabolism. To explore if this upregulation of FASN may contribute to increased histone crotonylation and cardiomyocyte hypertrophy, FASN expression was targeted using siRNA or the pharmacological inhibitor TVB-2640. Inhibition of FASN expression or activity reduced the ET-1-dependent increase in H3K18cr, as well as a trend toward increased total histone Kcr (Fig. 2C, D). Moreover, targeting FASN also attenuated ET-1 induced expression of ANP, consistent with reduced cellular hypertrophy. This was substantiated by measuring cardiomyocyte cell size, which was elevated after ET-1 treatment but attenuated upon FASN inhibition by TVB-2640 (Fig. 2E). These findings suggest that FASN may be an important therapeutic target to limit cardiomyocyte hypertrophy.

### Downstream activation of mTOR and p300 leads to elevated histone crotonylation

The mechanism behind FASN upregulation following hypertrophic stimulation was found to be linked to mTOR activity. The inhibition of mTOR by rapamycin attenuated the upregulation of FASN and H3K18cr by ET-1 (Fig. 3A). Furthermore, we identified a reciprocal mode of FASN and mTOR activation. Here inhibition of FASN led to loss in mTOR activation following hypertrophic stimulation (Fig. 3B). This suggests that activity of both enzymes is tightly coupled with inhibition of FASN leading to suppression of prohypertrophic mTOR signaling. As mTOR is known to enhance p300 activity^20^, we hypothesized that this mechanism may underlie the observed increase in histone Kcr during cardiomyocyte hypertrophy and loss with rapamycin treatment. To investigate whether p300 may contribute to the increase in histone crotonylation, we assessed its acetylation and crotonylation following hypertrophic stimulation. Here p300 Kcr was found to be increased following ET-1 treatment that was attenuated by rapamycin (Fig. 3C). Also, there was a switch from p300 acetylation to crotonylation after ET-1 treatment, thus suggesting also a change in p300 substrate utilization.

**Figure 3.**
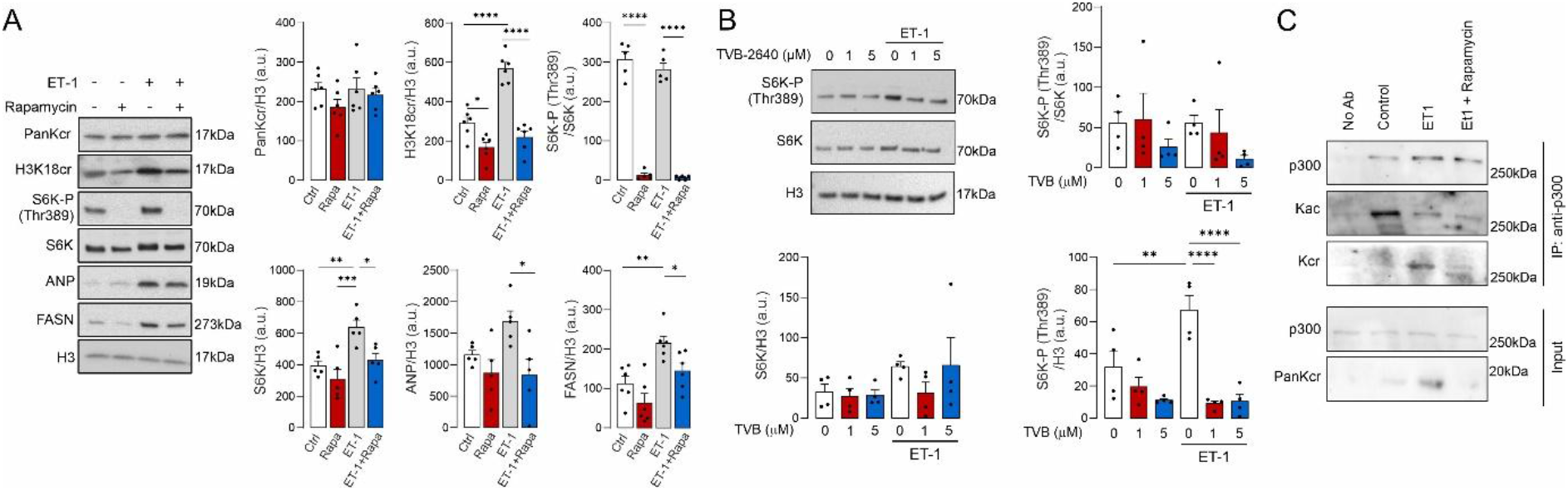
FASN dependent activation of mTOR leads to ET-1 dependent elevation in histone crotonylation. **A,** Histone Kcr, H3K18Cr, FASN, S6K, S6K-P and ANP expression in cardiomyocytes treated with or without rapamycin and in the presence or absence of ET-1 (n=5-6). **B**, S6K phosphorylation in cardiomyocytes treated with or without the FASN inhibitor TVB-2640 and in the presence or absence of ET-1 (n=4). **C**, p300 Kac and Kcr detected after affinity capture of p300. *P<0.05; **P<0.01; ***P<0.001; ****P<0.0001. Comparisons were made using 1-way ANOVA followed by the Tukey post hoc test.

### FASN-dependent hypertrophic gene expression is associated with increased histone H3K18 crotonylation

To establish the role of FASN upregulation following hypertrophic stimulation we assessed the impact of this enzyme on gene expression during cardiomyocyte hypertrophy. Here we compared differential gene expression between control and ET-1-treated cells with that of ET-1 versus ET-1 with TVB-2640. Pathway analysis revealed that FASN drove expression of genes associated with cardiomyopathy and cardiomyocyte hypertrophy (Fig. 4A-B). These included genes encoding motor proteins and those involved in ECM-receptor interaction that are implicated in cardiomyocyte hypertrophy (Fig. 4C)^21,22^. In complementary studies targeting FASN using siRNA also revealed similar findings where FASN was implicated in driving gene expression associated with cardiomyocyte hypertrophy (Fig. S3A, B).

**Figure 4.**
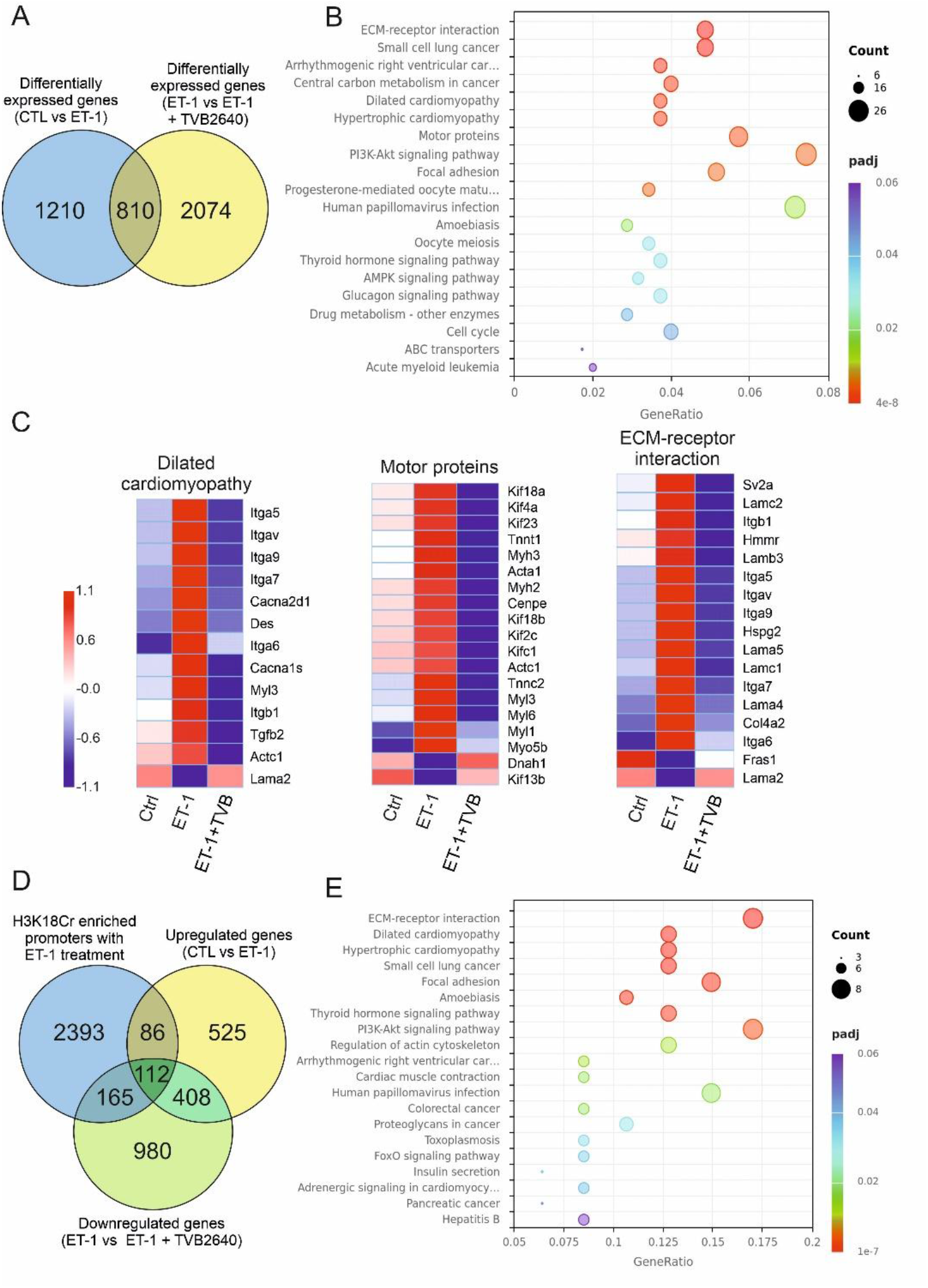
FASN and downstream elevation in histone crotonylation drives expression of hypertrophic genes. **A**, Venn diagram of genes upregulated by ET-1 treatment with those that are downregulated with ET-1 and FASN inhibitor TVB-2640 (n=4). **B**, Pathway analysis of common genes that are upregulated by ET-1 treatment and those that are downregulated with ET-1 and FASN inhibitor TVB-2640. **C**, Heat maps for pathways associated with dilated cardiomyopathy, motor proteins and ECM-receptor interactions identified from the pathway analysis (n=4). **D**, Venn diagram of genes upregulated by ET-1 treatment (RNA-seq, n=4), downregulated with ET-1 and FASN inhibitor TVB-2640 (RNA-seq, n=4) and those enriched for H3K18cr in hypertrophic cardiomyocytes (Cut&Run, n=3). **E**, Pathway analysis of common genes that are upregulated by ET-1 treatment (RNA-seq), downregulated with ET-1 and FASN inhibitor TVB-2640 (RNA-seq) and those enriched for H3K18cr in hypertrophic cardiomyocytes.

To determine whether FASN mediated expression of genes associated with hypertrophy were mediated by increased histone crotonylation, we performed CUT&RUN sequencing to map genomic regions with elevated H3K18cr. Integration of promoter-associated H3K18cr enrichment with FASN-regulated genes during cardiomyocyte hypertrophy enabled identification of pathways likely influenced by this epigenetic modification (Fig.4D).

Subsequent pathway analysis revealed that H3K18cr is associated with FASN-dependent expression of genes implicated in cardiomyopathy and cardiomyocyte hypertrophy (Fig. 4E). Therefore, it is likely that FASN-dependent elevation of histone Kcr contributes to cardiomyocyte hypertrophy.

### FASN expression correlates with hypertrophic gene expression in human HCM

To investigate whether FASN expression is associated with hypertrophic transcriptional programmes in human disease, we analysed an existing RNA-seq dataset generated from cardiac tissue of patients with HCM^13^. Expression of each gene was correlated with FASN expression across patient samples (Fig. 5A). This analysis revealed strong positive associations between FASN expression and the canonical hypertrophic marker genes NPPA and NPPB (Fig. 5B, C). We next compared the strength of association between FASN and HCM-related genes with that observed for background genes (Fig. 5D). HCM-related genes showed significantly stronger correlations with FASN than background genes. FASN expression also showed a statistically significant, although modest, association with sample-wide mean gene expression (Fig. S4A). To assess whether this contributed to the observed relationships, expression values of every gene were residualised against sample-wide mean expression (Fig. S4B). Following residualisation, the associations between FASN and NPPA/NPPB were retained (Fig. S4C), as well as those between FASN and HCM-related genes (Fig. S4D). These analyses support an association between FASN expression and disease-related hypertrophic gene expression in human HCM.

**Figure 5.**
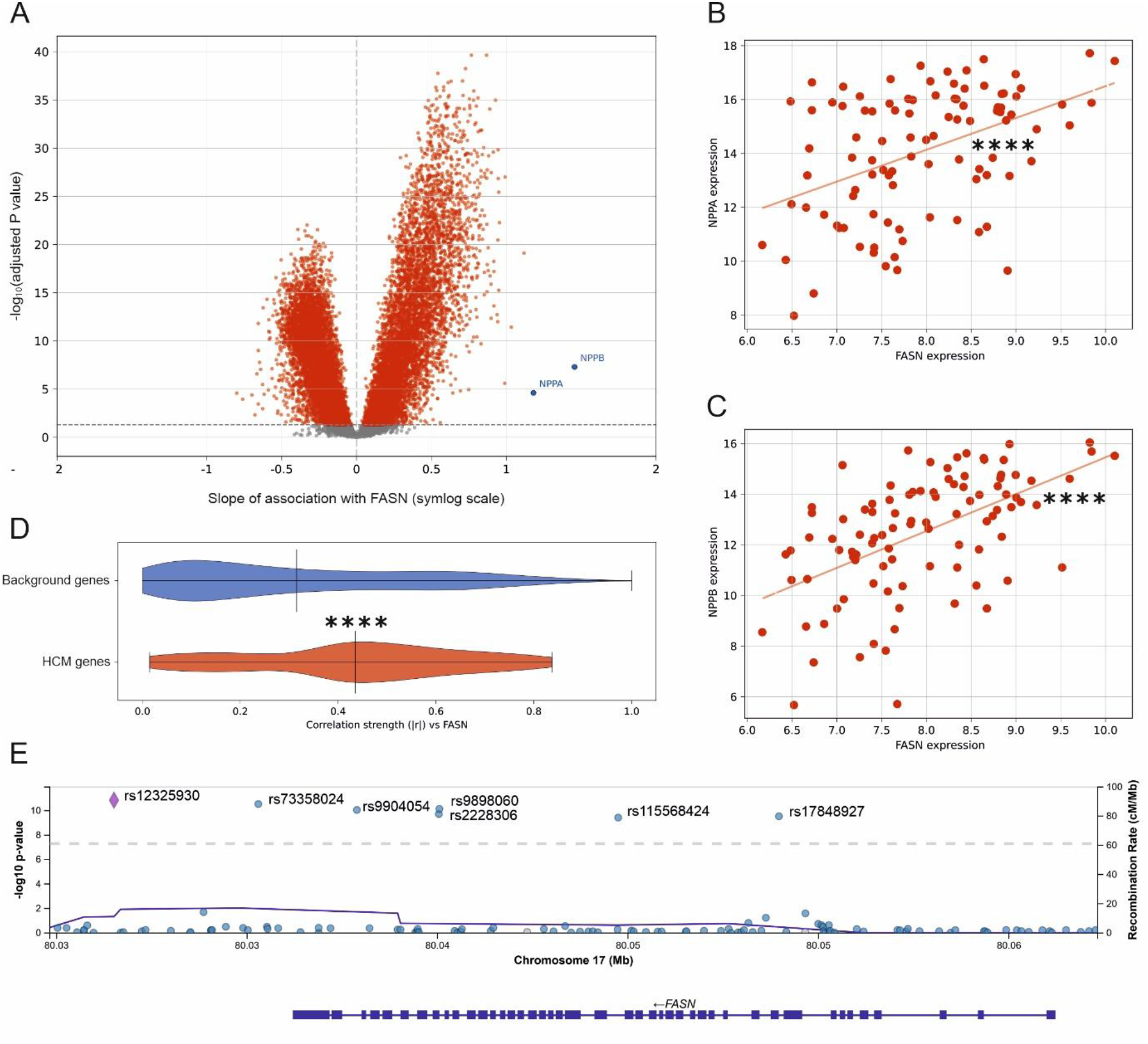
FASN expression correlates with markers of hypertrophy in patients with hypertrophic cardiomyopathy. **A**, Volcano plot showing regression slope and statistical significance from gene-wise association with FASN in RNAseq of cardiac tissue from HCM patients. **B**, Scatter plot comparing FASN expression with NPPA expression. **C**, Scatter plot comparing FASN expression with NPPB expression. **D**, Violin plot comparing correlation coefficient |r| with FASN between HCM-related genes and background genes. **E,** Regional association (LocusZoom) plot showing the association of genetic variants within the FASN locus with heart failure recorded as the underlying (primary) cause of death (ICD-10 code I50.9)^15^. ****P<0.0001. Comparisons were made using ordinary least-squares analysis (**A**, **B**, **C**) or Welch’s t-test (**D**).

To investigate whether genetic variants associated with heart-failure-related mortality are also linked to FASN expression, we examined publicly available expression quantitative trait locus (eQTL) datasets. The variant rs12325930, located within the *FASN* locus, which is the lead variant most significantly associated at genome-wide significance level with heart failure as the underlying cause of death (ICD-10 code I50.9, Fig 5E), showed evidence of an association with FASN expression during cardiomyocyte differentiation in the dataset reported by Elorbany et al (β = 1.069, SE = 0.381, P = 0.00535)^16^. The positive interaction coefficient indicates that this variant is also associated with enhanced FASN expression.

## Discussion

Here, we identify a previously unrecognized signaling axis that drives cardiomyocyte hypertrophy through FASN-dependent upregulation of histone crotonylation. Previous studies have shown that histone crotonylation is elevated in human HCM and in mice treated with AngII, with histone H3K18cr identified as a key modification site^9^. Consistent with these findings, we observed increased H3K18cr in cardiomyocytes exposed to hypertrophic stimuli, as well as a pronounced increase in total histone crotonylation in the hypertrophic myocardium seven days following transverse aortic constriction (TAC). Together, these observations provide a strong rationale for histone crotonylation as an important epigenetic regulator in the development of cardiomyocyte hypertrophy, likely through enhancement of gene expression^5^. This concept is further supported by evidence demonstrating that histone crotonylation is a potent driver of transcription, including some evidence that crotonate treatment promotes enrichment of NFATc3 at the Nppb promoter^9^. Given the strong correlation between histone crotonylation and cardiomyocyte hypertrophy, we sought to define the underlying biochemical mechanisms and to establish the functional importance of this epigenetic modification in regulating cell growth.

We first evaluated the potential role of enoyl-CoA hydratase short chain 1 (ECHS1), as its loss has previously been linked to increased histone crotonylation under hypertrophic conditions. However, we did not observe a reduction in ECHS1 expression when histone crotonylation was elevated. In contrast, knockdown of ECHS1 resulted in a decrease in histone H3K18cr levels. Although this finding contradicts earlier reports, ECHS1 is involved in the generation of crotonyl-CoA. This is supported by studies where ECHS1 knockdown in oocytes was found to suppress histone crotonylation^18^, and in human stem cells ECHS1 was found to act as a crotonyl-CoA synthase, contributing to histone Kcr^19^.

Given that ECHS1 was unlikely to account for the increase in histone crotonylation observed during hypertrophy, we investigated alternative pathways. This led to the identification of FASN as a key mediator in the upregulation of histone crotonylation. FASN expression was increased following hypertrophic stimulation of cardiomyocytes and after TAC surgery.

These findings are consistent with previous reports demonstrating elevated FASN expression in mouse models of heart failure and in the myocardium of patients with class IV cardiomyopathy^23^. Notably, FASN protein levels are reduced in patients following left ventricular assist device placement, indicating that FASN upregulation is reversible with mechanical unloading of the failing myocardium. Moreover, transgenic mice with cardiomyocyte-specific overexpression of FASN develop features of heart failure^24^. In agreement with these studies, we found that FASN upregulation not only enhanced histone crotonylation but also directly promoted cardiomyocyte hypertrophy. Furthermore, increased FASN activity following hypertrophic stimulation enhanced the expression of genes associated with dilated and HCM, as well as genes encoding motor proteins and components of extracellular matrix–receptor interactions implicated in cardiac hypertrophy^21,22^.

To further elucidate the underlying mechanism, we examined the interplay between FASN and mammalian target of rapamycin (mTOR) signaling. Activation of mTOR was required for the upregulation of both FASN expression and histone crotonylation following endothelin-1 (ET-1) treatment. This is consistent with the established role of mTOR in driving cardiac hypertrophy and promoting FASN expression^25,26^. In addition, inhibition of FASN also attenuated mTOR activation in response to hypertrophic stimulation, suggesting a tightly coupled regulatory relationship between FASN and mTOR. One possible explanation for the reciprocal regulation of FASN and mTORC1 is the accumulation of malonyl-CoA following FASN inhibition, which has previously been identified as an endogenous ATP-competitive inhibitor of mTORC1^27^. This process may also explain the observed increase in histone Kcr, as mTOR can directly activate p300^20^, a known histone crotonyltransferase^5,28^. Consistent with this, inhibition of mTOR resulted in a loss of H3K18cr. Moreover, p300 crotonylation was enhanced during hypertrophic stimulation, in line with increased crotonyltransferase activity, and this effect was attenuated by rapamycin. In contrast, p300 acetylation was reduced following hypertrophic stimulation^29^. Together, these findings suggest a shift in p300 substrate utilization during hypertrophy, from acetyl-CoA to crotonyl-CoA.

To establish the functional importance of histone crotonylation in mediating FASN-dependent cardiomyocyte hypertrophy, we identified genes likely to be regulated by histone H3K18cr during hypertrophic stimulation. These genes were predominantly associated with dilated and HCM, as well as extracellular matrix–receptor interaction pathways^21^, further supporting a central role for histone crotonylation in pathological cardiac growth.

To assess the translational relevance of our findings, we analysed cardiac RNA-seq data from patients with obstructive HCM and found that FASN expression was positively associated with NPPA and NPPB, two canonical markers of pathological hypertrophy. We also identified a variant located within the FASN locus that associated with heart-failure-related mortality and also enhanced FASN expression in cardiomyocytes. This further supports the potential link between FASN expression and heart failure progression in humans. However further studies are required to determine whether variants that enhance FASN expression do indeed couple to the development or progression of heart failure.

Together, our findings support a model in which FASN coordinates mTOR signaling and histone crotonylation during cardiac hypertrophy. Although further in vivo validation is required, this pathway may represent a therapeutically tractable mechanism for limiting pathological cardiac remodeling and progression towards heart failure.

## Data availability

All data that support the findings of this study are available from the corresponding author upon request. The RNA-seq and CUT&RUN data have been deposited in the NCBI GEO database (GSE343751).

